# Voltage-gated sodium channels in neocortical pyramidal neurons display Cole-Moore activation kinetics

**DOI:** 10.1101/164210

**Authors:** Mara Almog, Tal Barkai, Angelika Lampert, Alon Korngreen

## Abstract

Exploring the properties of action potentials is a crucial step towards a better understanding of the computational properties of single neurons and neural networks. The voltage-gated sodium channel is a key player in action potential generation. A comprehensive grasp of the gating mechanism of this channel can shed light on the biophysics of action potential generation. Most models of voltage-gated sodium channels assume it obeys a concerted Hodgkin and Huxley kinetic gating scheme. Here we performed high resolution voltage-clamp experiments from nucleated patches extracted from the soma of layer 5 (L5) cortical pyramidal neurons in rat brain slices. We show that the gating mechanism does not follow traditional Hodgkin and Huxley kinetics and that much of the channel voltage-dependence is probably due to rapid closed-closed transitions that lead to substantial onset latency reminiscent of the Cole-Moore effect observed in voltage-gated potassium conductances. This may have key implications for the role of sodium channels in synaptic integration and action potential generation.

## Introduction

The action potential is the fundamental unit of communication between neurons. Thus, a detailed mechanistic understanding of the action potential is a paramount goal in neuroscience. One major milestone dates back more than sixty years when Hodgkin and Huxley (1952) showed that action potential generation required a fine balance between the fluxes of sodium and potassium ions through voltage-gated conductances. From then onwards, their kinetic model has served as a cornerstone for understanding the generation and propagation of action potentials in many systems, leading to enormous progress in accounting for the neuronal input/output transformation.

It thus comes as no surprise that the vast majority of simulations in neuronal physiology use the Hodgkin and Huxley formalism to simulate ion channel kinetics. However, a much more detailed picture of the mechanisms underlying membrane excitation has emerged over the years that challenges several features of the Hodgkin and Huxley model. For example, the classical Hodgkin and Huxley paradigm does not consider interactions between varying kinetic states, and in particular between activation and inactivation (Armstrong and Bezanilla 1977; Bezanilla and Armstrong 1977; Raman and Bean 2001; 1997). In contrast to physiologically oriented modeling, structure function investigations of voltage-gated channels usually describe channel kinetics using Markov chain models (Sakmann and Neher 1995). These structure function studies have shown, for almost all voltage-gated ion channels, that Hodgkin and Huxley-like kinetics are only a rough approximation of the kinetic behavior of the channels (Patlak 1991).

Two prime deviations from the classical Hodgkin and Huxley kinetics have been observed in neurons from the rodent CNS. The voltage-gated channels in many CNS neurons display prolonged, also known as persistent, currents (Astman et al. 2006; Brown et al. 1994; Crill 1996; Fleidervish and Gutnick 1996; Khaliq et al. 2006; Magistretti and Alonso 2002; Magistretti et al. 1999; Schwindt and Crill 1995; 1980). Persistent sodium currents have been observed in layer 5 pyramidal neurons as of the early days of whole-cell recordings (Crill 1996; Schwindt and Crill 1980; Stafstrom et al. 1982; 1984a; Stafstrom et al. 1984b) and together with resurgent voltage-gated sodium currents greatly increase the contribution of sodium conductance to the action potential (Astman et al. 2006; Brumberg et al. 2000; Fleidervish et al. 1996; Fleidervish and Gutnick 1996; Kole et al. 2008; Kole and Stuart 2008; Lewis and Raman 2014; Raman and Bean 2001; 1997; Raman et al. 1997; Stuart and Sakmann 1995). A number of studies have reported that the sodium channel activates with a longer delay than is predicted by the Hodgkin and Huxley model (Martina and Jonas 1997; Schmidt-Hieber and Bischofberger 2010). This delayed activation has been incorporated into the Hodgkin and Huxley formalism by adding a simple delay to the current onset (Martina and Jonas 1997) or incorporated into a Markov model of the channel (Schmidt-Hieber and Bischofberger 2010). Interestingly, one study, focusing on voltage-gated sodium channel activation at temperatures colder than room temperature reported that the channels activate without delay (Baranauskas and Martina 2006).

It is thus clear that the kinetics of voltage-gated sodium channels deviate considerably from the classical Hodgkin and Huxley model. However, there is no consensus how to best describe their kinetics. Clearly, the level of detail required by the model is greater when the aim is to correlate the structure and function of the channel than when the goal is to simulate the action potential (Almog and Korngreen 2016; Lampert and Korngreen 2014). To address these questions, we conducted voltage-clamp experiments primarily from nucleated patches extracted from the soma of layer 5 (L5) pyramidal neurons. We show that the gating mechanism does not follow traditional Hodgkin and Huxley kinetics. Specifically, much of the channel voltage-dependence is due to rapid closed-closed transitions leading to substantial onset latency reminiscent of the Cole-Moore effect observed in voltage-gated potassium conductances (Cole and Moore 1960).

## Methods

### Animals

All procedures were approved and supervised by the Institutional Animal Care and Use Committee and were in accordance with the National Institutes of Health Guide for the Care and Use of Laboratory Animals and the University’s Guidelines for the Use and Care of Laboratory Animals in Research. This study was approved by the National Committee for Experiments in Laboratory Animals at the Ministry of Health.

### Slice preparation

Slices (sagittal, 300 μm thick) were prepared from the somatosensory cortex of 12-16 days old Wistar rats that were killed by rapid decapitation using previously described techniques (Stuart et al. 1993). Slices were perfused throughout the experiment with an oxygenated artificial cerebrospinal fluid (ACSF) containing (mM): 125 NaCl, 25 NaHCO_3_, 2.5 KCl, 1.25 NaH_2_ PO_4_, 1 MgCl_2_, 2 CaCl_2_, 25 Glucose, 0.5 Ascorbate (pH 7.4 with 5% CO_2_, 310 mosmol/kg). All experiments were carried out at room temperature (25°C). Pyramidal neurons from L5 in the somatosensory cortex were visually identified using infrared differential interference contrast (IR-DIC) videomicroscopy (Stuart et al. 1993).

### Electrophysiology

Nucleated outside-out patches (Almog and Korngreen 2009; Korngreen and Sakmann 2000; Sather et al. 1992) were extracted from the soma of L5 pyramidal neurons. Briefly, negative pressure (180-230 mbar) was applied when recording in the whole cell configuration, and the pipette was slowly retracted. Provided that the retraction was gentle it was possible to obtain large patches of membrane engulfing the nucleus of the neuron. After the extraction of the patch, the pressure was reduced to 30-40 mbar for the duration of the experiment. All measurements from nucleated patches were carried out with the Axopatch-700B amplifier (Axon Instruments, Foster City, CA) using a sampling frequency of 100 kHz and filtered at 20 kHz. The capacitive compensation circuit of the amplifier was used to reduce capacitate transients. Nucleated patches were held at -60 mV. Leak was subtracted using a P/4 online protocol. Patch pipettes (4-7 MΩ) were coated with Sylgard (DOW Corning). To record sodium currents in nucleated patches the pipettes were filled with a solution containing 120 mM Cs-gloconate, 20 mM CsCl, 10 mM HEPES, 4 mM MgATP, 10 mM phosphocreatine, 1 mM EGTA, 0.3 mM GTP (pH=7.2, CsOH). The liquid junction potential of -11 mV generated by this pipette solution was not corrected for during the experiments or offline analysis. Cell attached recordings were performed from the soma of L5 pyramidal neurons using standard procedures (Hamill et al. 1981; Stuart et al. 1993). To record sodium currents in the cell attached mode the pipettes were filled with a solution containing 130 mM NaCl, 3 mM KCl, 2 mM MgCl_2_, 2 mM CaCl_2_, 10 mM HEPES, 4 mM TEA, 1 mM 4-aminopyridine, and 10 mM glucose (pH=7.3 with HCl). Cell attached patches were held at -20 mV relative to the membrane potential. Similar to the nucleated patches, the pipettes were coated with Sylgard and the leak was subtracted using a P/4 online protocol. All cell-attached recordings were performed using the AxoPatch-200B amplifier (Axon Instruments, Foster City, CA) using a sampling frequency of 100 kHz and filtered at 20 kHz.

### Data analysis and numerical simulations

All off-line data analyses including curve fitting were carried out with IGOR (WaveMetrics, Lake Oswego, USA). Experimental results were observed in cells from two or more animals. Therefore, all the results from a specific protocol were pooled and displayed as means± S.E.M. Simulations of ionic currents were programmed using NEURON 7.3 (Hines and Carnevale 1997). All simulations were performed with an integration interval of 5 μs to ensure a stable numerical solution of the differential equations. Ion channel models were implemented using the NMODL extension of NEURON (Hines and Carnevale 2000). The models of channels reported in the literature were downloaded from ModelDB (https://senselab.med.yale.edu/ModelDB/) and used without any changes to the code.

## Results

Sodium currents were recorded from nucleated patches after blocking voltage-gated potassium currents as detailed in the Methods. Fast inward currents could easily be detected following a depolarizing voltage clamp step from a holding potential of –110 mV (Fig. 1A). We only analyzed currents from which the linear leak was cleanly subtracted by the P/4 protocol, thus leaving no residual of the capacitance transient. This was quite straightforward when the capacitance transient did not saturate the amplifier or exceed the dynamic range of the A/D converter. Due to the small membrane surface the total capacitance of a nucleated patch was small and allowed for excellent temporal control of the voltage-clamp amplifier (Baranauskas and Martina 2006; Martina and Jonas 1997). We verified this by measuring the decay of the uncompensated capacitance transients that decayed to baseline in under 40 μs. To be on the safe side, we removed the first 50 μs from the data before analysis. Note that this may result in an under-estimation of the channel kinetics.

**Figure 1:**
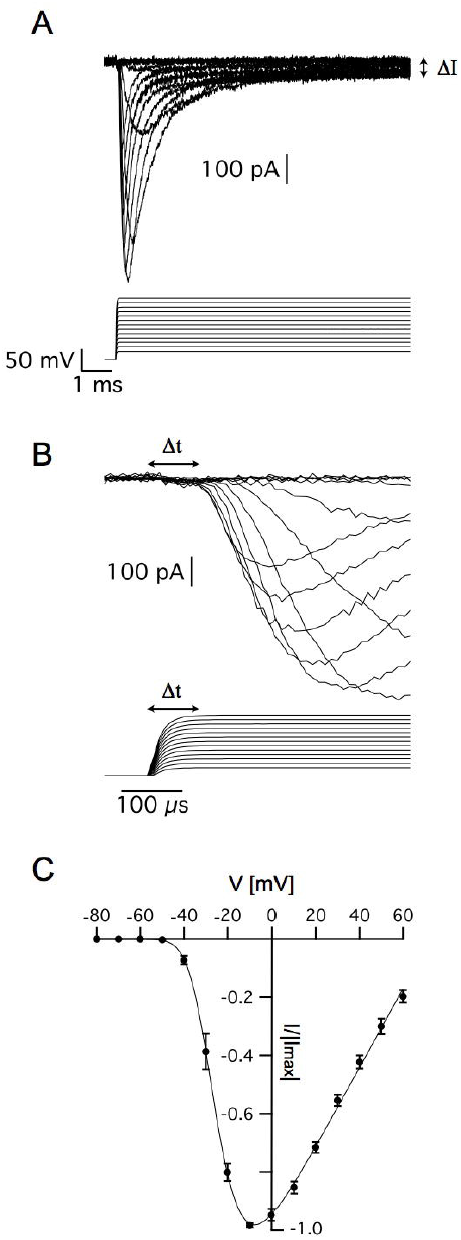
Voltage-gated sodium currents recorded in nucleated patches from cortical pyramidal neurons. A, representative recording of sodium currents from layer 5 neocortical pyramidal neurons. B, magnification of the time immediately following the voltage change showing the delayed activation of the sodium current. The vertical scale of the command potential is identical to the one presented in A. C, activation curve calculated from 9 nucleated patches. The current in each patch was normalized to the absolute value of the maximal current amplitude recorded at -10 mV to reduce variability between patches. The line connecting the points is the curve fit of Eqn. 1. Error bars are SEM.

As described elsewhere, the recorded currents displayed two features deviating from the classical description of voltage-gated sodium channels in that there was a sustained current (denoted by ΔI in Fig. 1A) and delayed activation (denoted by Δt in Fig. 1B). We first estimated the apparent voltage-dependence of channel activation by measuring the maximal current change in 9 patches. To reduce inter-patch variability, we normalized each patch to the maximal current obtained in this analysis. Plotting the maximal normalized current from 9 patches as a function of voltage produced a typical activation curve of voltage-gated sodium channels (Fig. 1C). Fitting this curve with

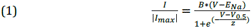

where *B* is a proportion constant, *E*_*na*_ is the sodium reversal potential, *V*_*0.5*_ is the voltage of half activation, and *z* is the slope of the Boltzmann curve, we obtained V_0.5_ =-24± 2 mV, z=5.2± 0.7 mV, and E_Na_ =75± 5 mV (n=9).

In the current study, we focused on the analysis of the delay in channel activation. The activation curve (Fig. 1C) suggested that above -10 mV, the channel would approach its maximal open probability. Thus, in the simple scheme for the activation (disregarding inactivation) of the channel suggested below it was reasonable to assume that for depolarized potentials all backward rate constants would be much smaller than the forward rate constants and could thus be neglected.

Scheme 1:

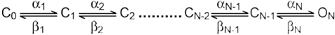

In this voltage range, the average delay between the onset of the voltage command and reaching the open state is given by;

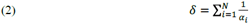

Assuming that the forward rate constants increase exponentially with voltage it is simple to show that δ will decrease as a function of voltage. Clearly, d is dependent δthe number of transitions leading to channel opening and the overall rate of the activation process. It is less sensitive to interdependence between rate constants in the activation scheme. The expression for δ in a sequential model only differs from a concerted model by a proportionality constant. Together, the number of transitions and rate of activation defined the amount of delay relative to the overall rate of activation or sigmoidicity of the activation process (Schoppa and Sigworth 1998a; b; Zagotta et al. 1994a; Zagotta et al. 1994b). In voltage-gated potassium channels the sigmoidicity was estimated by fitting an exponential function to the slowest part of the current trace followed by extrapolation to a zero current providing an estimate for δ (Schoppa and Sigworth 1998a; Zagotta et al. 1994b). In the current study, this was obviously not possible due to channel inactivation. Thus, we assumed that the initial stages of the activation were only marginally affected by inactivation. This assumption was put to the test at a later stage (see Figure 4). Thus, we only analyzed the current up to 50% of its maximal current by fitting it with

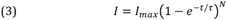

where τis the time constant and N is the apparent number of transitions in the Markov chain model (Fig. 2A). Eqn. 3 is not the general solution to scheme 1 but rather the concerted version of that model. Using it considerably reduced the number of free parameters to be fitted. Previously it was shown that fitting this equation produces practically indistinguishable results from the general solution to scheme 1 (Schoppa and Sigworth 1998a; Zagotta et al. 1994b) due to the difficulty of distinguishing between certain kinetic models using whole-cell currents (Milescu et al. 2005).

**Figure 2:**
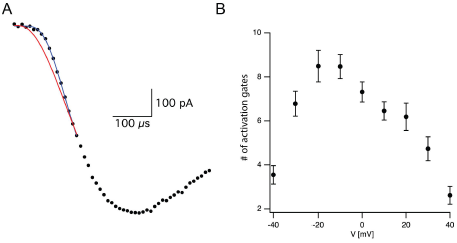
Estimating activation sigmiodicity for the voltage-gated sodium channel. A, Representative curve fit of Eqn. 3 to the onset of current activation (black line). The current was recorded after a voltage step from -110 mV to -10 mV. The red line is a fit of the same data to Eqn. 3 in which the number of activation gates was set to 3 as in the Hodgkin and Huxley model. The black line is the fit of Eqn. 3 using the number of activation gates as a free parameter. This specific fit generated a value of N=9± 1. B, The apparent number of activation gates, obtained by fitting Eqn. 3 to data from 9 patches, is plotted as a function of the membrane potential. In all cases the patch was held at -110 mV for 150 ms prior to stepping to the test potential. Error bars are SEM.

Fitting eqn. 3 to the current onset with N=3 as in the Hodgkin and Huxley model (red line in Fig. 2A) failed to provide a complete fit for the data. Fitting the current onset this time using N as a free parameter produced an improved fit with N=9.1 for the trace displayed in Figure 2A (black line in Fig. 2A). As predicted by Eqn. 2, the apparent number of closed states signifying the sigmoidicity of the current trace decreased as a function of voltage above -10 mV (Fig. 2B). We extended the fit to lower potentials that displayed smaller sigmoidicity probably due to the participation of backward transitions slowing the current trace. Thus, our phenomenological analysis supported a model for the activation of voltage-gated sodium channels in L5 pyramidal neurons containing considerably more closed states than the three gates assumed by the Hodgkin and Huxley model.

Early results for the voltage-gated potassium channel of the giant squid axon indicated that holding potential could modulate latency before the onset of activation (Cole and Moore 1960). Hyperpolarization of the membrane potential increased the latency of current activation. These authors reported that these effects increased the apparent number of activation gates in the potassium channel from 4 to 25. This may suggest that the prolonged delay and steep current activation we observed in our recordings from voltage-gated sodium channels were due to a “Cole-Moore effect”.

We tested this prediction by varying the holding potential prior to channel activation. Using a standard protocol designed for measuring the steady-state inactivation of voltage-gated sodium channels we varied the holding potential from -110 mV to -25 mV for a period of 150 ms prior to stepping the potential to zero mV to record the current (Fig. 3A). The maximal current change following each pre-pulse was normalized to the maximal current change recorded following the pre-pulse to -110 mV. The resulting normalized inactivation curve, averaged across patches, is presented in Figure 3B. Fitting this curve to a Bolzmann function gave V_0.5_ =-64.5± 0.5 mV with z=8.3±0.3 mV (n=11). Next, we fitted the first 50% rise of the current to Eqn. 3. As predicted, for a sequential activation model containing several closed states, the apparent number of activation gates decreased steadily as the pre-pulse potential increased (Fig. 3C) which is consistent with results obtained for the sodium channel in the squid giant axon (Keynes and Rojas 1976). This supports the hypothesis that the activation of voltage-gated sodium channels in L5 pyramidal neurons displayed Cole-Moore kinetics, probably due to many sequential closed transitions before the open state. However, at this stage we could not dismiss the possibility that these results might be an experimental artefact related to the extraction of the nucleated outside-out patch. Thus, we also recorded sodium currents in the cell-attached mode of the patch-clamp technique and subjected them to the same analysis. Out of the 78 patches only three had large enough currents for the analysis of current onset. Estimating the apparent number of activation gates from these patches (Fig. 3D) showed a similar dependence of the apparent number of gates on the command potential to that obtained from nucleated patches (Fig. 3C).

**Figure 3:**
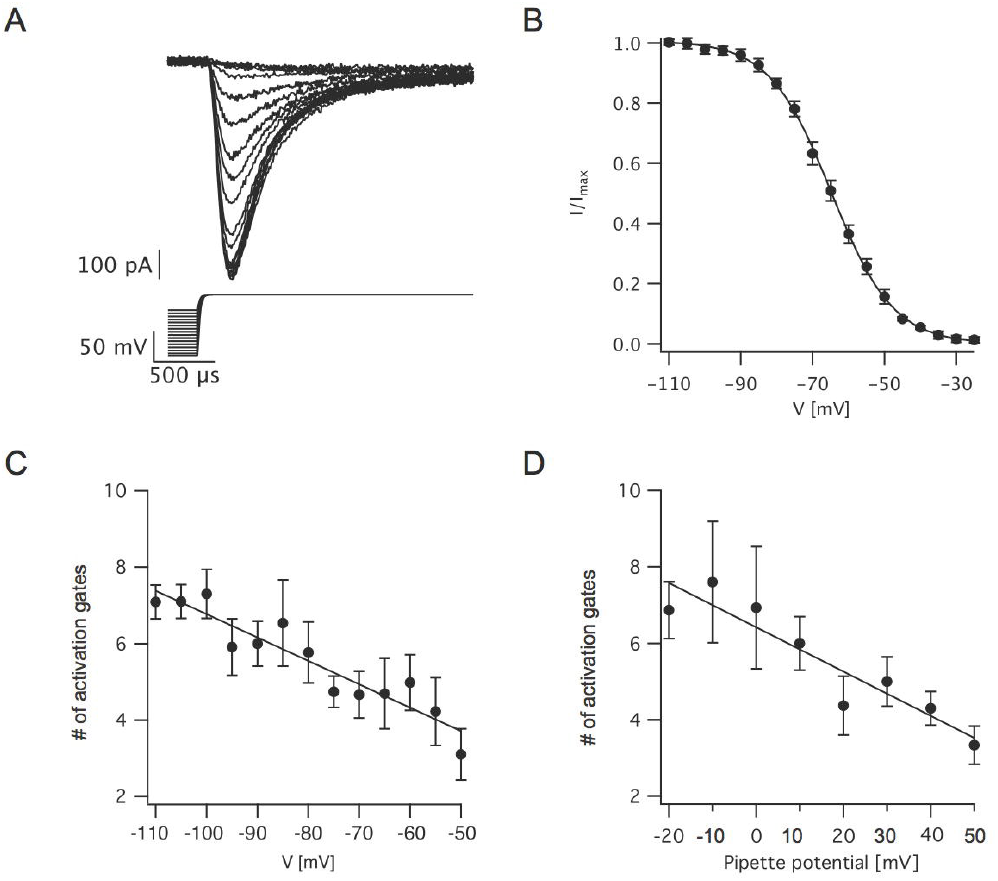
Estimating the voltage-dependence of activation sigmoidicity in the channel inactivation experiment. A, representative recording of currents evoked by a voltage-clamp protocol designed to measure steady-state inactivation. The 150 ms pre-pulse to various potentials was truncated to facilitate the display of the current. In all trials, the voltage was stepped to 0 mV following the pre-pulse. B, Steady-state inactivation curve calculated from 11 recordings identical to those displayed in A. The current was normalized to the trace obtained following a pre-pulse to -110 mV. The smooth line of a fit of a Boltzmann function. C, The apparent number of activation gates, obtained by fitting Eqn. 3 to data from 9 nucleated patches, plotted as a function of the pre-pulse potential. D. The apparent number of activation gates, obtained by fitting Eqn. 3 to data from 3 cell-attached patches, plotted as a function of the pre-pulse potential which is displayed as the pipette potential since the true membrane potential is unknown. Error bars in are SEM in all subfigures.

The analysis of sigmoidicity using Eqn. 3 has never been carried out on currents from voltage-gated sodium channels. To test this procedure, we applied it to three known models of voltage-gated sodium channels (Fig. 4). It was simple to predict that the sigmoidicity of the Hodgkin and Huxley (Hodgkin and Huxley 1952) model (Fig. 4Ai) would be constant and equal to three. In fact, throughout the voltage range of channel activation our analysis of the initial current onset produced this expected result (Fig. 4Aii). Fitting Eqn. 3 to the current onset of the inactivation protocol produced the expected result (N=3) whereas the pre-pulse potential was below the activation threshold of the conductance (Fig. 4Aiii). Above-activation threshold lower values of N were obtained.

**Figure 4:**
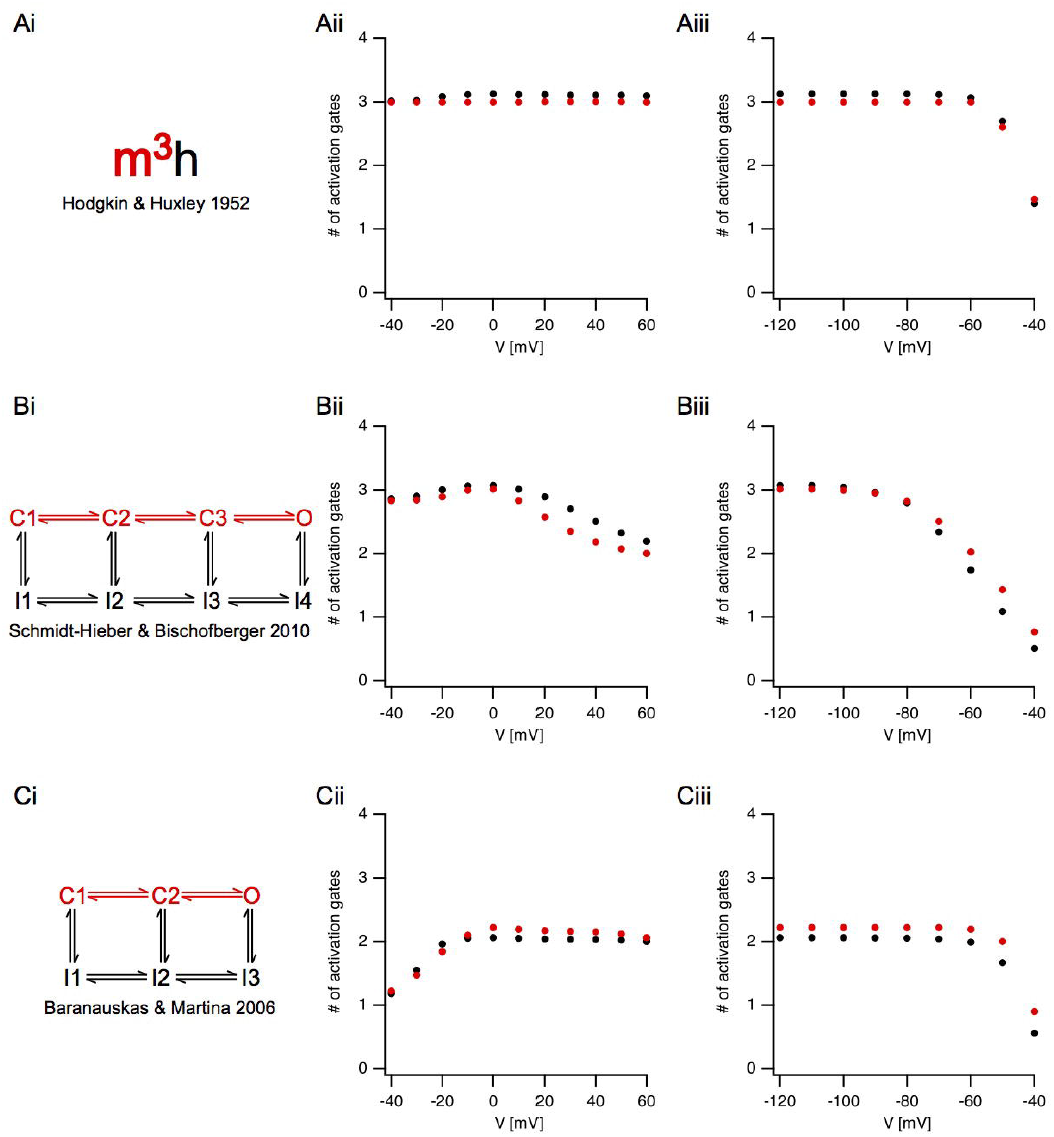
Estimating activation sigmoidicity in models of voltage-gated sodium channel. Ai, Bi, Ci, A schematic representation of the model. The pertinent part of the model for activation appears in red. The source of each mode is indicated below each scheme. Aii, Bii, Cii, The apparent number of activation gates, estimated by fitting Eqn. 3 to the initial onset of the current, plotted as a function of the potential in an activation protocol (the pre-pulse was 150 ms to -100 mV followed by a step to the test potential). The black circles are the values obtained from currents simulated using the full model. The red circles are the values obtained from currents simulated using only the activation part of the model. Aiii, Biii, Ciii, The apparent number of activation gates, estimated by fitting Eqn. 3 to the initial onset of the current, plotted as a function of the potential in the inactivation protocol (the pre-pulse varied from -100 mV in steps of 10 mV followed by a step to the 0 mV). Color coding is identical to that of Aii, Bii, and Cii.

Fitting Eqn. 3 up to 50% of the maximal current amplitude was performed under the assumption that the fit was not distorted by inactivation kinetics. To test this assumption, we removed the inactivation from the Hodgkin and Huxley model, and simulated non-inactivating currents. Fitting the onset of these currents to Eqn. 3 produced almost identical results to those obtained when fitting the full model (red markers in Fig. 4). This verified that our assumption regarding the negligible effect of inactivation on the onset analysis was justified.

Next, we applied the onset analysis to two Markov models of voltage-gated sodium channels from the central nervous system. The model proposed by Schmidt-Hieber and Bischofberger (2010) for hippocampal granule cells consists of three closed states (Fig. 4Bi). We thus predicted that the onset analysis for some of the traces would be N=3. This turned out to be the case for both the activation (Fig. 4Bii) and inactivation (Fig. 4Biii). However, unlike the Hodgkin and Huxley model, the number of gates predicted for the Schmidt-Hieber and Bischofberger model varied as a function of voltage as the occupancy of the states in the Markov model shifted from the first closed state all the way to the open state. This is very clear from Figure 4Biii that depicts the dependence of the sigmoidicity on the pre-pulse potential. As the pre-potential increased, the channel occupied closed states closer to the open state. Upon activation, the current trace activated with a shorter delay, thus leading to a smaller value of N when fitted to Eqn. 3. Consistent with the Schmidt-Hieber and Bischofberger model, fitting the Baranauskas and Martina (2006) model of cortical neurons (Fig. 4Ci) to Eqn. 3 provided an approximate value of N=2 for most potentials,. Deviations from this value either in the activation (Fig. 4Cii) or inactivation (Fig. 4Ciii) protocols could easily be accounted for by the relative occupancy of the model in the closed states along with the rate of transition from the last closed state to the open state. To further test this assumption regarding the negligible influence of the inactivation kinetics on the curve fit we deleted the inactivation from the two Markov models and repeated the analysis only using the linear model containing closed states leading to the open state (marked in red in Figs. 4Bi and 4Ci). These simulations generated values for N that were very similar to the full models (red symbols in Fig. 4) further justifying the assumption we made in the analysis of the experimental traces (Figs. 2 and 3). Adding a tail of several closed states to either one of these Markov models increased the value of sigmoidicity in positive correlation with the number of added closed states (simulations not presented).

## Discussion

Here we showed that the activation of voltage-gated sodium channels in layer 5 pyramidal neurons from the rat neocortex does not follow traditional Hodgkin and Huxley kinetics. In addition, much of the channel voltage-dependence could be attributed to rapid closed-closed transitions that led to substantial onset latency reminiscent of the Cole-Moore effect observed in voltage-gated potassium conductances (Cole and Moore 1960). Thus, the activation of the channels was state dependent and displayed a distinct delay if activated from hyperpolarized potentials, whereas there was almost immediate activation if activated from potentials closer to the action potential threshold.

The consequences of this state dependent activation to action potential firing are straightforward. We observed that the sigmoidicity of the sodium current increased as the holding potential was more negative (Fig. 3). It is important to note that the increase in sigmoidicity was correlated to the increase in the delay before onset of channel activation. Given that the resting membrane potential of L5 pyramidal neurons ranged roughly from -70 mV to -60 mV, our crude analysis predicted that stepping from these potentials to above the channel activation threshold should generate a current displaying kinetics obeying ~ 4 activation gates. However, the physiological changes to the membrane potential are slower than the standard voltage-clamp step since they are limited by the membrane time constant. Thus, during synaptic integration changes to closed state occupancy of the sodium channel will clearly be faster than membrane potential changes. Consequently, as the membrane potential approaches the action potential threshold the sodium channel is likely to activate almost without delay.

The Cole and Moore (1960) study of the voltage-gated potassium channel of the giant squid axon showed that the holding potential could modulate the latency before the onset of activation. Hyperpolarization of the membrane potential increased the latency and the steepness of the current during activation (Cole and Moore 1960). They reported that these effects increased the apparent number of activation gates in the potassium channel from 4 to 25. This “Cole-Moore” effect has also been observed in the shaker potassium channel (Hoshi et al. 1994; Zagotta et al. 1994a; Zagotta et al. 1994b), the large conductance calcium activated potassium channel (Horrigan and Aldrich 1999; Horrigan et al. 1999), and in the voltage-gated sodium channel in the squid giant axon (Keynes and Rojas 1976). More recently, delayed activation has been observed in voltage-gated sodium channels in the rat central nervous system (Baranauskas and Martina 2006; Martina and Jonas 1997; Schmidt-Hieber and Bischofberger 2010).

Of the studies reporting an activation delay in the rat central nervous system, two stand out since they present somewhat conflicting results. Baranauskas and Matina (Baranauskas and Martina 2006) recorded voltage-gated current from several types of neurons in the rat CNS. They reported, in apparent contradiction to our results, that voltage-gated channels activate without a delay. Contrary to this study, Schmidt-Hieber and Bischofberger (2010) reported that voltage-gated sodium channels display a distinct, voltage-dependent, activation delay in hippocampal granule cells.

The discrepancy between these studies can be reconciled in view of the data we present here. Most of the recordings made by Baranauskas and Matina were carried out from a holding potential of -70 mV. Conversely, Schmidt-Hieber and Bischofberger applied a 50 ms pre-pulse to -120 mV to remove inactivation. This difference in the initial conditions of the experiments changes the occupancy of the closed states of the channels. Holding at -70 mV may be regarded as a more physiological initial condition which causes the channel to occupy a closed state close to the open state. The downside of this holding potential is of course the considerable inactivation that reduces the total current. The model presented by Baranauskas and Matina (Fig. 4Ci), in which a single transition from the closed to the open state determines the activation kinetics of sodium currents, clearly captures some of the activation kinetics of the voltage-gated channel. The model presented by Schmidt-Hieber and Martina (Fig. 4Bi) that considers the voltage-dependence of the initial delay, contains an additional closed state. As the membrane potential approaches threshold, this closed state is largely unoccupied, which makes this model more similar to the Baranauskas and Matina model.

The role of the mechanism proposed here in action potential generation in L5 pyramidal neurons is probably part of a more complex mechanism. The action potential is known to be generated at the distal end of the axon initial segment (Hu et al. 2009; Palmer and Stuart 2006; Popovic et al. 2011), though under some conditions it is generated at the first Ranvier node. That said, several studies have clearly shown complex crosstalk between the initiation zone and the soma (Foust et al. 2010; Khaliq and Raman 2006; Palmer and Stuart 2006; Telenczuk et al. 2017). The complexity of the axon initial segment derives from the composition and distribution of its voltage-gated ion channels, especially the voltage-gated sodium ion channels. Recently it was shown that the patch-clamp technique is probably inappropriate for determining the density of ion channels in the axon initial segment because of the ways in which these channels are anchored in the cytoskeleton with Ankyrin-G, which suggests, as models of the action potential have predicted, that the sodium channel density is high in the axon initial segment (Kole et al. 2008). Converging findings from labeling studies (Lorincz and Nusser 2008; Royeck et al. 2008) and sodium imaging (Baranauskas et al. 2013; Fleidervish et al. 2010; Palmer and Stuart 2006) suggest that the sodium channel density in the axon initial segment may be substantially larger than the soma, roughly ~ 4,000 pS/μm2. Using electrophysiology and immunohistochemistry, it was elegantly demonstrated that the axon initial segment has two different voltage-gated sodium channels, Nav1.2 and Nav1.6, and that these channels have graded distributions along the membrane (Hu et al. 2009); Nav1.6 has a higher density in the distal part of the initial segment, and this anatomical complexity explains much of the axon initial segment’s physiological complexity. In the current study, we recorded from nucleated patches extracted from the soma. Thus, our conclusions regarding the role of the delayed activation of the voltage-gated sodium channel, while logical, should be probed by recording the activity of NaV1.6 in axonal patches or expression systems.

Regardless of the role played by the activation delay in action potential firing, our results clearly indicate a need to re-evaluate the application of the Hodgkin and Huxley formalism to modeling the role of ions in neuronal excitability and the value of replacing it with Markov chain models. Like all models, the Hodgkin and Huxley model approximates reality. As such, it performs well under certain conditions and fails in others. As noted above (Cole and Moore 1960; Moore 2015), Hodgkin and Huxley carried out most of their recordings from a holding potential of -70 mV. Thus, it is not surprising that they obtained a relatively short delay, leading to the claim of three activation gates for the sodium channels and four activation gates for the potassium channels. However, this selection of activation gates eliminated a dynamic aspect of the channel from the model. This can clearly be seen in Figure 4Aii in which the sigmoidicity does not change as a function of voltage. The large sigmoidicity we report here indicates that the full model for the voltage-gated channel should probably contain at least 7-8 closed states. The full scope of this model, which should also include the fast and slow inactivation processes, has yet to be determined.

## Acknowledgments

This work was supported by a grant from the German-Israel Foundation to AK and AL (#1091-27.1/2010).

